# Leveraging spatial scale and temporal variation to optimize estimates of invasive spread rates

**DOI:** 10.1101/2025.02.06.636321

**Authors:** Joseph Keller, Sebastiano De Bona, Matthew R. Helmus

**Affiliations:** Integrative Ecology Lab, Center for Biodiversity, Department of Biology, Temple University, 1925 N. 12th Street, Philadelphia, PA 19122; Department of Biology, Bryn Mawr College, Bryn Mawr, Pennsylvania, USA

**Keywords:** alpha hull boundary, grape pest, integrated pest management, invasion curve, invasion front delineation, invasive species, jump dispersal, *Lycorma delicatula*, range limit size, spatial scale, spread rate

## Abstract

Estimating the extent and speed of an invasive species’ spread is crucial for optimizing surveys and management, but this estimation is challenging due to the scale-dependent nature of spread. For instance, when discretizing occurrence data to grids to estimate the extent of invaded ranges, larger cells can decrease boundary precision, while smaller cells can increase the risk of missing invaded areas. Moreover, on shorter timescales such as year-to-year comparisons, spread rates can exhibit lags, accelerations, and slowdowns. We present a multiscale spread optimization methodology developed for the spotted lanternfly (*Lycorma delicatula*), a pest impacting grapes that is spreading westward across the continental United States. We analyzed a dataset of >900,000 occurrence records covering spread from this pest’s initial detection in eastern Pennsylvania in 2014 to its invasion front in Chicago, IL, in 2023. First, we delineated the annual U.S. invasion front using square grids with varying cell sizes and α-convex hulls with different boundary resolutions. For each method and scale, we regressed invaded range area against year using both simple linear and logistic nonlinear models. Coarser spatial scales yielded faster estimated spread rates, and logistic models outperformed linear regressions, indicating a lag, acceleration, and slowdown in annual spread rate. Next, to determine the spatial scale that best captured the invasion front boundary, we used cross-validation, optimizing the F_β_ metric, which balances recall and precision. Emphasizing recall generally favored coarser spatial scales, and α- convex hulls outperformed grid-based methods in delineating invasion fronts. Finally, *L. delicatula* has dispersed long distances from its primary epicenter, establishing satellite populations. We delineated each satellite population each year since its inception using optimized α-hull values and measured their year-to-year boundary expansion. Like the primary epicenter, these satellite populations showed accelerating expansion, at least until merging with the primary invasion. Overall, the expansion rate of the primary invasion has slowed since 2017, decreasing from a maximum of 21 km/year, possibly due in part to control efforts, although the exact cause remains unresolved. Considering spatial scale and temporal variation can optimize the analysis of invasive spread and motivate early interventions to manage nascent epicenters before their spread accelerates.

## Introduction

The spread patterns of invasive species strongly influence their economic impact (Epanchin-Niell & Liebhold, 2015). Detailed knowledge of how far and fast the targeted outbreak is spreading can guide the choice of the most appropriate management objectives, which can range from eradication, to slowing spread, to limiting impacts in invaded areas (Rejmanek & Pitcairn, 2002; Liebhold & Kean, 2019). Additionally, accurate knowledge of spread rates can improve the specific strategies deployed to achieve those objectives, guiding optimal spatial and temporal allocation of surveillance and management efforts (Andrew & Ustin, 2010; Epanchin-Niell & Hastings, 2010; Triska & Renton, 2018).

Temporal variation complicates the process of characterizing the spread patterns of invasive species. Theory predicts that invasions’ spread rates should generally follow a predictable pattern over time (Shigesada et al., 1995). Initially, spread is expected to be slow, as populations take time to grow; lag phases between initial introduction and subsequent rapid spread are in fact frequently observed (Coutts et al., 2018; Crooks & Soulé, 1999; Memmott et al., 2005). After this lag phase, spread accelerates and may approach a constant asymptotic rate for some time, depending on the specific characteristics of the invasion (Hastings et al., 2004). Finally, the spread rate in the invaded range must slow as more and more of the potentially suitable range becomes occupied (e.g. Ward et al., 2020). As the invaded range becomes saturated, dispersing propagules are more likely to end up in areas that are already occupied or unsuitable for survival, resulting in no further spread. The emerald ash borer (*Agrilus planipennis*), for example, has had differing spread rates in the U.S. over time, with breakpoint analysis revealing 4 distinct spread periods that followed the expected slow-rapid-slow pattern closely (Ward et al., 2020). The spread rate of the spongy moth (*Lymantria dispar*) in North America has also varied over time, though less predictably, with rapid spread between 1900 and 1915, followed by a slower spread phase from 1916 to 1965, and another rapid spread phase between 1966 and 1990 (Liebhold et al., 1992). Managers benefit from a detailed understanding of spread rates variation and from information on the status of the targeted invasion.

The need to select an appropriate spatial scale for analysis complicates efforts to characterize invasive species’ spread patterns because methods for characterizing these patterns rely on delineating invaded areas from uninvaded areas, a process inherently dependent on scale. Estimation of the size of the invaded range and the location of the invasion front depend on the spatial scale used in the analysis. For example, invasion fronts are drawn based on geopolitical units (Tobin et al., 2007), trap locations (Tobin et al., 2013), point locations of reported presences (Wallace et al., 2020), remote sensed data (Lonsdale, 1993), and expert opinion (Storer, 1937).

Regardless of the approach, at coarser scales, species can be said to occupy entire nations, while at much finer scales the spatial footprint of each individual will make up the species’ distribution. Scaling must be considered whether the invaded range is evaluated based on areal measures (grid cells or geopolitical units occupied) or boundary drawing (where the accepted level of concavity and the inclusion/exclusion of small outlying populations and/or unoccupied pockets depends on scaling parameters). With the growing availability of geolocated point records describing the occurrence patterns of organisms, identifying an appropriate scale for analyzing such data is becoming increasingly important (Shirey et al., 2021).

Given these issues of spatial scale and temporal variation, researchers have used various methods to measure spread patterns. For example, a common approach estimates spread rate by tracking changes in invaded area over time, approximating the area as a circle and calculating annual changes in its radius using square-root area regression. (Hastings et al., 2004; Skellam, 1951). Alternatively, spread rate may be estimated by measuring the distances between invasion fronts from consecutive time steps. Vectors between boundaries may be measured radiating from a central point (typically the origin of the invasion) (Tobin et al., 2007), oriented perpendicular to the invasion front, or drawn as the shortest distance between points (Tisseuil et al., 2016). In general, without a clear consensus on which method is best to estimate spread patterns, authors often apply multiple approaches to the same spatio-temporal datasets (e.g. Liang et al., 2019, Ward et al., 2020, Cook et al., 2021).

Well-documented invasions provide valuable case studies to test and compare methods for estimating spread patterns (Hastings, 1996). The spotted lanternfly (*Lycorma delicatula*) is an invasive planthopper whose spread in the United States has been thoroughly surveyed through the combined efforts of state and federal agencies (De Bona et al., 2023). A univoltine phloem feeder, the spotted lanternfly damages vineyards by feeding on grapevines, and has spread rapidly in the U.S. since its first detection in Berks County in southeastern Pennsylvania in 2014 (Cook et al., 2021; Urban & Leach, 2023). Satellite populations are common because this pest hitchhikes on a variety of transported materials and readily establishes (Huron et al., 2022). Here, we present a detailed analysis of the spread patterns *L. delicatula* has exhibited thus far in the United States, focusing on the following questions:

1. How has spread rate varied over time?
2. How does spatial scale affect spread rate estimates?
3. How do different methodologies for delineating invaded ranges affect spread rate estimates?
4. What spatial-scale parameters result in best-fitting delineations based on cross validation?
5. Do satellite populations follow similar patterns of spread as the primary invasion?

## Methods

To address our research questions, we evaluated change in the area of the invaded range using both grids and α-convex hulls and assessed evidence for temporal variation in spread rates by comparing the fit of simple linear regressions and logistic curves. We identified best-performing spatial scales for analysis using cross validation and, based on those parameters, measured spread rates around satellite population epicenters that have arisen over time. These results provide an updated understanding of this pest’s spread in North America.

### Data

To characterize the growing distribution of *L. delicatula* in the United States, we gathered presence and absence records documenting the sampling effort from state agencies and the USDA covering the period from 2014, when the pest was first found, through 2023 (Table 1).

**Table 1:** Sampling effort has varied over time as the *L. delicatula* invasion in the U.S. continued to grow. Numbers of presence and absence records for *L. delicatula* from government (state and federal) sources over the biological years 2014 through 2023 are listed, along with two measures of the invasion size: the invaded area and the length of the boundary, both as assessed with an α- convex hull with α’ = 35 km. Biological years turn over on March 31/April 1. To show how sampling effort compared to the size of the invasion, we report the number of observations per km^2^ area and per km of boundary perimeter.

| Biological Year | Number of presences | Number of presences with confirmed establishment | Number of absences | Invaded area (km<sup>2</sup>) | Invasion boundary length (km) | Sampling effort (Obs. per km<sup>2</sup> area) | Sampling effort (Obs. per km boundary) |
| --- | --- | --- | --- | --- | --- | --- | --- |
| 2014 | 62 | 26 | 308 | 33.5 | 26.3 | 10.6 | 14.1 |
| 2015 | 2,876 | 2,479 | 4,800 | 274.0 | 77.3 | 28.0 | 99.4 |
| 2016 | 2,780 | 2,286 | 6,482 | 1718.1 | 188.3 | 5.4 | 49.2 |
| 2017 | 1,333 | 895 | 7,843 | 7200.7 | 403.5 | 1.3 | 22.7 |
| 2018 | 4,705 | 3,937 | 95,552 | 16387.6 | 644.1 | 6.1 | 155.7 |
| 2019 | 10,441 | 7,477 | 120,535 | 29708.3 | 1080.5 | 4.4 | 121.2 |
| 2020 | 10,867 | 3,765 | 65,985 | 39980.6 | 1688.2 | 1.9 | 45.5 |
| 2021 | 10,799 | 5,116 | 53,944 | 54924.1 | 2535.9 | 1.2 | 25.5 |
| 2022 | 12,593 | 6,812 | 52,853 | 83808.6 | 3487.0 | 0.8 | 18.8 |
| 2023 | 8148 | 5,951 | 32,803 | 112695.3 | 3989.9 | 0.4 | 10.3 |

All analyses were based on biological years reflecting the annual life cycle of *L. delicatula*. Each biological year’s records reflect the presence or absence of individuals that hatched in that year and the eggs they produced. Egg masses laid in one calendar year but discovered before hatch in the next, for example, would be included in the first year’s data. In practice, the biological year turns over on 30 April or 1 May, depending on the year. The anonymized data are available at https://github.com/ieco-lab/lydemapr and a full description of processing to compile the data is detailed in De Bona et al. (2023).

### Grid-based boundaries

To distinguish invaded areas from uninvaded areas, we divided the eastern U.S. where the pest has spread into grids of varying scales and assessed the presence of *L. delicatula* within each cell. In each year, cells where at least one established *L. delicatula* population was noted in that year or in any prior year were considered invaded, while cells without any of these records were considered uninvaded. While we used regular grids with varying resolution here, this process is closely analogous to methods based on geopolitical units such as townships, counties, states, and countries (Liang et al., 2020).

### α convex hull boundaries

To test an alternative method for delineating invaded and uninvaded areas, we fit α- convex hulls. α-convex hulls are mathematically well-defined (Edelsbrunner et al., 1983) and are capable of accurately locating range boundaries for irregularly shaped and disjointed ranges (Burgman & Fox, 2003). This approach to locating species range boundaries has been applied most frequently in investigations of the range size of threatened species (IUCN Standards and Petitions Committee, 2024), though it has also been used previously to estimate range sizes of potential invasives (Hui et al., 2011) and to track the spread of biological invasions (Urban et al., 2008). α-convex hulls are constructed using triangulation of presence points, with a single parameter (α) determining the resolution of the boundary, which consists of arcs of open circles with a radius equal to α (Pateiro-Lopez & Rodriguez-Casal, 2022). When α is very small, each presence is included as a disjointed point, and when α is infinitely large, the α-convex hull is congruent with the minimal convex hull (Edelsbrunner et al., 1983). For intermediate α values, polygons with varying resolution are generated. We used the *ahull* function from the *alphahull* package (Pateiro-Lopez & Rodriguez-Casal, 2022) and the *edmaps* package (Camac et al., 2021) in R (R Core Team, 2023) to fit α-convex hulls and process hulls.

To aid comparison with the grid-based analysis, we evaluated α values that resulted in circles with area equal to that of the square grid cells we evaluated. To do this, we report scaling results using a derived parameter α’, which is simply the edge length of a square with equal area to the circle with radius α, calculated as follows:

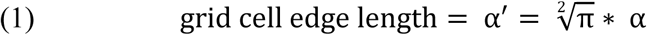

For our analyses, we evaluated spread patterns produced using cells with grid edge lengths and α’ values of 1, 2, 4, 6, 8, 10, 12, 14, 16, 18, 20, 25, 30, 35, 40, 45, 50, 75, and 100 km (19 in total).

We additionally included α’ = 1000 km to approximate the minimum convex hull. In a few cases, hull fitting failed due to linearity of points in small clusters. In these cases, we jittered points randomly up to 10 m away in both northing and easting and refit the hull.

### Square-root area-time regression

We employed square-root area-time regression to estimate *L. delicatula* spread rate in the U.S. We used the *st_area* function from the *sf* package in R to measure the area of the invaded range as determined by each boundary drawing method and parameter setting (Pebesma et al., 2023). Then, we calculated the effective range radius (ERR) (Shigesada et al., 1995):

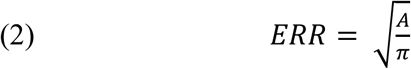

where *A* is the invaded area. ERR is the distance from the center to the edge of a circle with equivalent area to the invasion. For each bounding method and scale, we fit simple linear models regressing the effective range radius against year. We also fit logistic nonlinear curves following the form

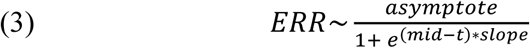

where *t* is the year, *asymptote* represent the eventual upper limit, *mid* represents the time at which growth is the fastest, and *slope* is a parameter controlling the slope of the curve. We used the *nls* and *SSlogis* functions in the *stats* package in R (R Core Team, 2023). The slope of the linear model is the estimated spread rate. For the logistic curve, spread rate varies over time, and is estimated by tangents to the curve. We evaluated models’ fit to the data based on AICc, the small sample corrected Akaike Information Criterion (Burnham & Anderson, 2002).

### Cross validation to select optimal scale for invaded range delineation

Both the grid-based and the α-convex hull methods for bounding the invaded range require the user to select a scale parameter. For grid-based bounding, the user selects the length of the cells’ edge. For α-convex hulls, the user chooses parameter α, which controls the boundary’s resolution. We tested 19 and 20 parameter values, respectively (listed in the previous section), with these two methods, and used cross validation to identify which parameter values resulted in boundaries that most effectively distinguished locations of *L. delicatula* presence from locations of absence. First, we randomly split the dataset into 5 subsets. We then performed cross validation separately for grid-based bounding and α-convex hulls (*methods*), for each year from 2014 to 2023 (*focal year*), and for each of the tested parameter values (*parameter*). For each combination of method, focal year, and parameter, we generated boundaries using a dataset that excluded one of the 5 subsets (the excluded subset later used as test data). Presence records from the year under investigation and from all prior years were used to create the boundary.

Then, we evaluated how effectively the resulting boundary captured presences and excluded absences by comparing it to the withheld test data (Appendix S1: Figure S1). We used the portion of the test data from focal year to evaluate both the precision (proportion of points included in the boundary that were truly presences) and recall (the fraction of all presence records that were included in the boundary). To balance the number of presences and absences used in evaluation, we resampled absences to ensure equal sample sizes. We note that imperfect detection may underlie some of the recorded absences, rather than lack of colonization.

Nonetheless, improving precision remains a valuable goal, even with potential failed-detection absences present in the data set.

We calculated recall and precision for each combination of method, focal year, and parameter, summing false positives, false negatives, and true positives across the 5 folds. Then for each method, year and scale, we calculated F*_β_* scores:

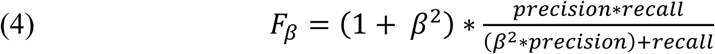

where *β* is a positive value that reflects the emphasis placed on recall, such that recall is *β* times as important as precision (Chinchor, 1992). The F*_β_* score provides a flexible metric for evaluating the scale-dependent delineations of invaded area from uninvaded area that reflects the decisions made by stakeholders (Hauser et al., 2016). This metric ranges from 0 to 1, with 1 reflecting perfect precision and recall. We evaluated scales’ performance using *β* values ranging from 1 (equal weight for presence and recall) to 4 (four times the emphasis placed on recall). Varying *β* reflects differing levels of importance that stakeholders may place on accurately capturing all presence points (high recall, emphasized with higher *β*) versus only indicating presence at specific locations where the targeted pest is found (high precision). Decision makers may determine the relative weight given each of these goals by selecting differing *β* values. We discuss alternative metrics in the discussion, but note that because bounding methods result in binary predictions rather than continuous values which could be split based on a threshold, we were unable to use metrics that depend on continuous scores such as area under the receiver operating curve (Lawson et al., 2014). We calculated F*_β_* scores for each year of observation, and with contingency table values summed across years, giving an estimate of the overall preferred values of cell edge length and α’.

Irregularity in the shape of the invaded range may vary with time and with spatial scale. To better understand the influence of the invaded range’s shape on the selection of a preferred scaling parameter, we calculated an index of circularity for each α-convex hull as the inverse of the normalized perimeter. Normalized perimeter is calculated by dividing a shape’s perimeter by the perimeter of a circle of equal area (Patton, 1975). This index of circularity ranges from 0 to 1, with low values indicating a shape differs dramatically from circular, while a value approaching 1 indicates similarity to a circle. Hulls were clipped to the coastline to remove any areas falling in the ocean prior to calculating this metric.

### Satellite population spread analysis

Multiple distinct satellite populations of *L. delicatula* have arisen in the U.S. after long-distance dispersal events. These distinct satellite populations provide additional cases where the early years of spread after introduction can be investigated. We fit α-convex hulls to the distribution of *L. delicatula* in each year, using the overall best performing value of α’ with β = 2 (35 km) and identified all distinct satellite populations by checking all polygons and seeing which overlapped each other in consecutive years. In cases where two or more formerly distinct populations merged, the oldest joining population was said to continue and any younger populations to no longer exist as distinct units. To focus on adequately sampled outlying populations, we excluded from analysis any outlying populations with fewer than 20 recorded absences within a 5 km buffer around their margins in each year. We measured each satellite population’s area in each year that it was present using the *st_area* function in the *sf* package (Pebesma et al., 2023).

We measured distances between consecutive years’ boundaries for all epicenters. While the α-convex hull provides a useful tool for bounding points where *L. delicatula* is established, it does not result in a “regular” border as described in Sharov et al. (1995), that is, a border where all lines radiating from an interior point cross the boundary only once. Islands, folds, and gaps in the boundary remain. For each cluster, we evenly spaced 300 points around the boundary for a given focal year, *t*, using the *st_line_sample* function (Pebesma et al., 2023). Then, for each of these points, we measured the shortest distance from that point to the border of any boundary polygon from the previous year*, t-1*, using the *st_distance* function. This approach accounted for instances when two or more disjunct populations merged from year to year. In these cases, the boundary displacement distance was measured to whichever of the two clusters was closer in the prior year.

## Results

The full database we built contained 259,723 records of *L. delicatula* establishment and 441,012 records of absence, including public records (Table 1). Established *L. delicatula* populations have been reported as far north as 43.21 °N, as far south as 35.63 °N, and as far west as 87.64 °W (Figure 1). The primary invasion has grown to cover large portions of Pennsylvania, New Jersey, Delaware, and Maryland. By 2023, far-flung outlying populations were established in several additional states, including Michigan, Illinois, Tennessee, North Carolina and Kentucky.

**Figure 1:**
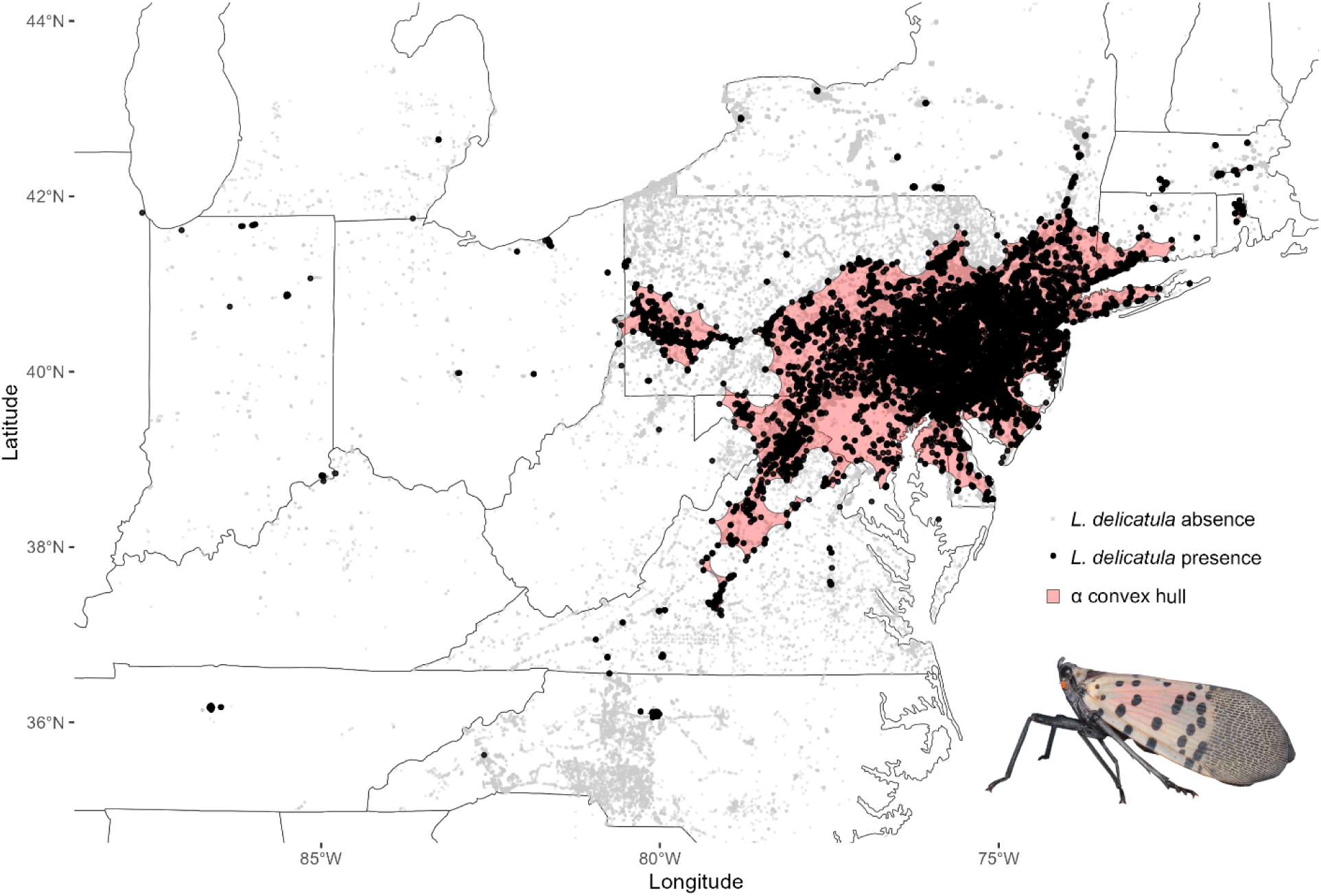
*Lycorma delicatula* (spotted lanternfly) occurrences in the United States as of 2023. Surveyed properties where government agencies have documented *L. delicatula* establishment are shown as black occurrence points, while those with no *L. delicatula* discovered are light grey. The invasive range (red polygon) was estimated as an α-convex hull (best-fit α’ = 35 km). See DeBona et al. (2023) for data and data aggregation methods.

### Square-root area-time regression

Spread rate estimates were substantially higher at coarser spatial scales (Figure 2, Appendix S1: Table S1-S4). This was true for both grid- and α-convex hull-based methods of bounding the invaded area. With coarser scaling parameters, *L. delicatula*’s invasion in the U.S. was estimated to cover a larger area, resulting in larger values for the spatial extent of the invasion (ERR) and more rapid increase in ERR over time (Figure 2). Generally, the fitted logistic curve had better fit to the data than the simple linear regression as measured by AICc (Figure 2, Appendix S1: Table S1-S4). The varying slope of the logistic curve precludes reporting a single spread rate, but the maximum spread rate offers a useful comparison across spatial scales, despite never being fully realized annually. Across the range of scales tested, the maximum spread rate increased from 32 km/year (2 km scale) to 218 km/year (50 km scale) for grids and 13 km/year to 116 km/year for α-convex hulls (Figure 2 E-F). Coarser scales also resulted in later estimated peak spread rate timing. For both grid and α-convex hulls, as scale increased, the peak shifted from occurring in 2018 to occurring in 2020

**Figure 2:**
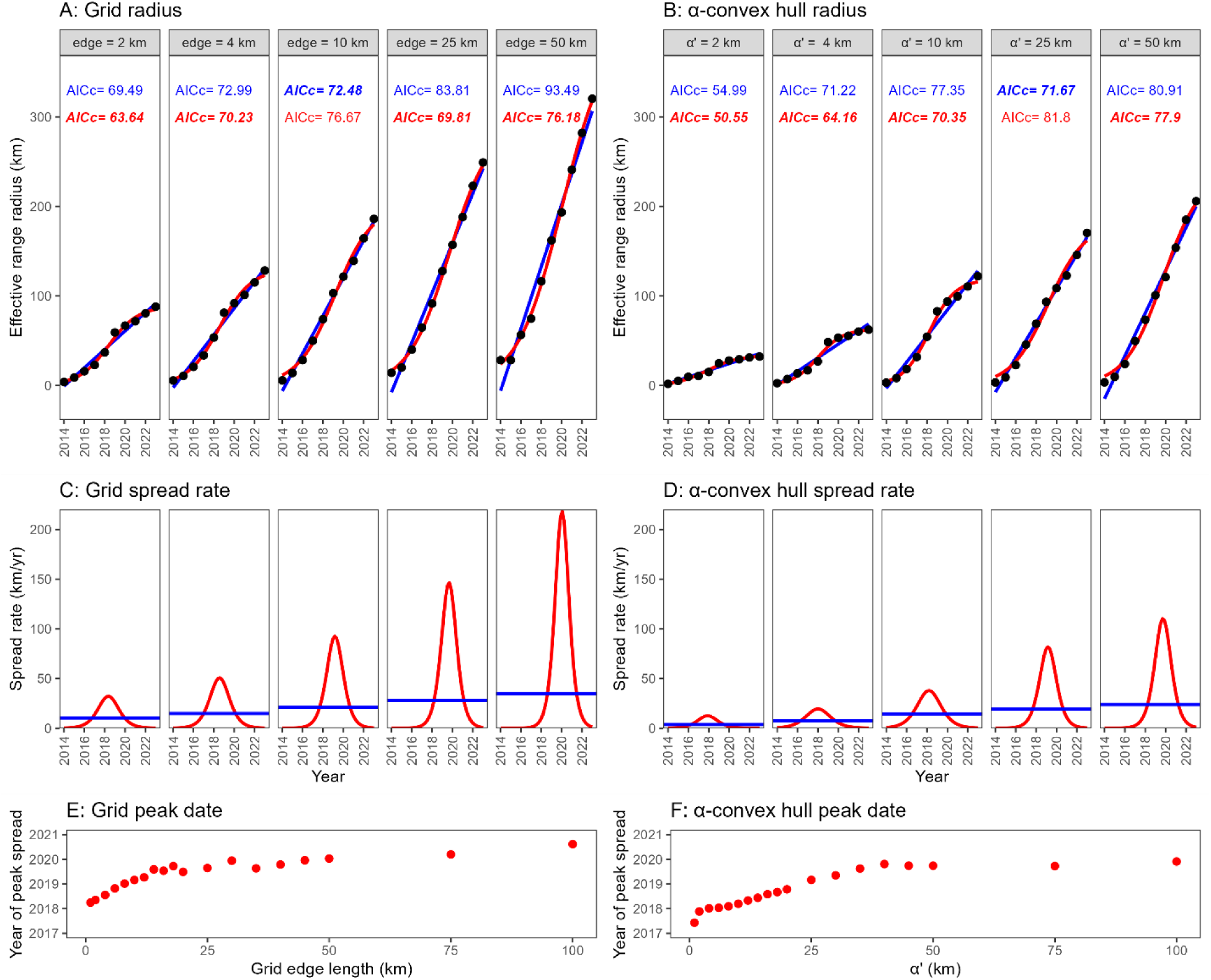
*Lycorma delicatula*’s estimated spread rate increased with spatial scale (grain size). Invaded range area was estimated based on grids: (A, C, E), for which grid cell edge length varied, and α-convex hulls (B, D, F), for which hull resolution varies (i.e., α’, eq. 1) Methods). To estimate spread rate from model slopes, we converted range area to effective range radius (radius of a circle of equal area) and fit simple linear regression models (blue) and logistic curve models (red), regressing ERR against year. We evaluated their fit using AICc (A, B). For most scales, logistic models fit better than linear models. Coarser scales of occurrence aggregation had higher spread rates (C, D), and later timing of peak spread as estimated from the logistic models (E,F), though the date of peak spread plateaued around 2020 as spatial coarseness increased.

### Cross validation to select optimal method scale for invaded range delineation

α-convex hulls generally delineated the location of the invasion front more accurately than grid-based boundaries, as measured by F_β_ score based on 5-fold cross validation. The best-performing α-convex hull fit the test data better than the best-performing grid-based boundaries for each year (Figure 3A). Differences between grid performance and α-convex hull performance were larger at small values of β. Due to the superior performance of α-convex hulls, we focus on this method for the remainder of cross validation results.

**Figure 3:**
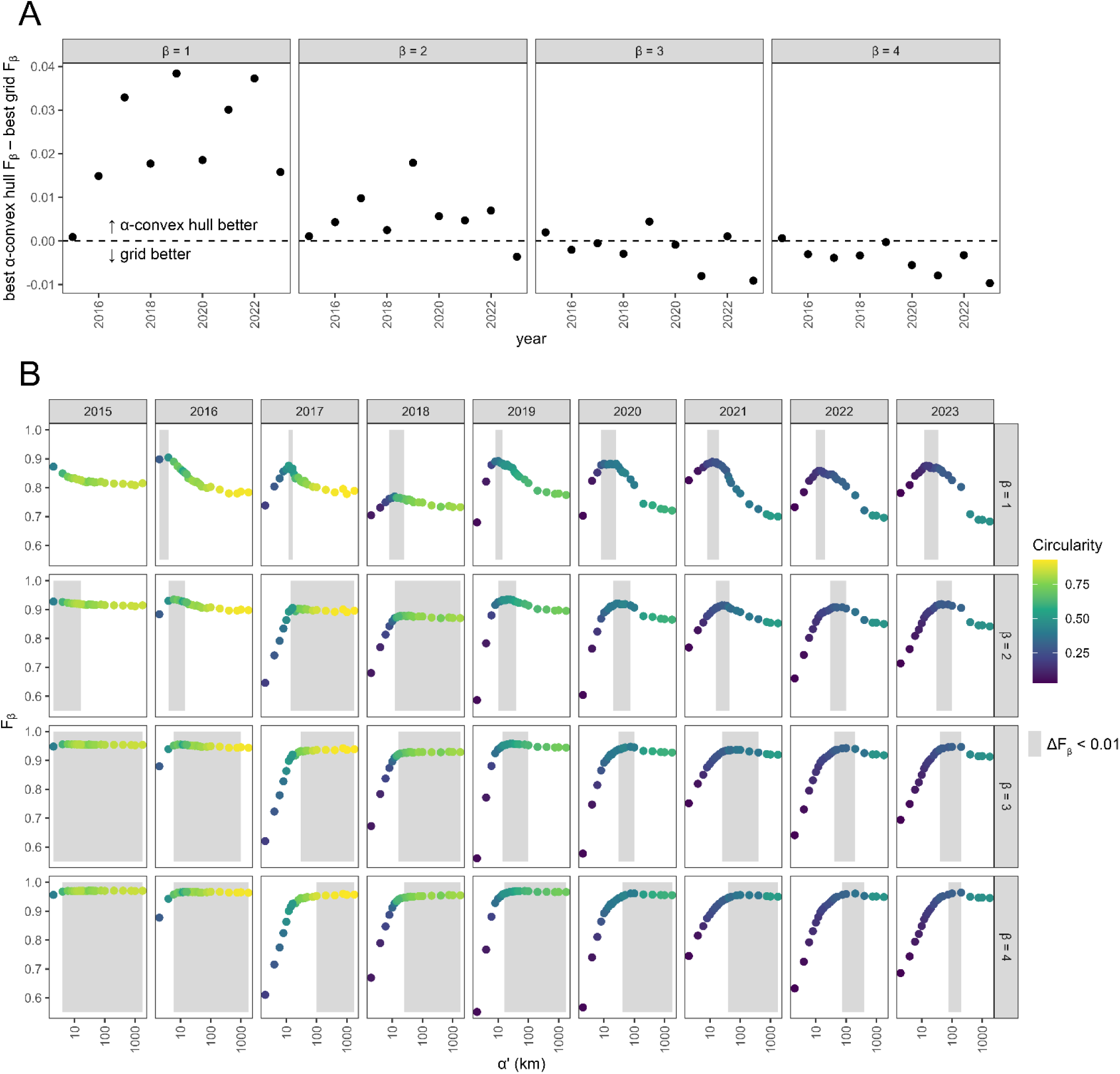
Best performing spatial scales varied over time and with the relative importance placed on recall versus precision. Using 5-fold cross-validation, we evaluated *L. delicatula* invasion boundaries in the U.S. annualy, comparing grids with varying cell-edge lengths to α-convex hulls with varying α’ resolution values (eq. 1). Boundaries were assessed using the F_β_ score, with β from 1 (equal weight on recall and precision) to 4 (quadruple weight on recall). α-convex hulls generally outperformed grids, except at high β where performace was comparable (A). The F_β_ scores for each year (columns, panel B), and β values (rows in panel B) were calculated, with the top-performing α’ values marked as shaded areas (within 0.01 of observed maximum) Best- performing α’ values increased with β (shaded areas shift right moving down columns) and when the invasion was more circular (yellow points), many values of α’ performed similarly, indicated by the wider shaded areas in the lower left panels.

The α value that performed best increased over time (Figure 3B, compare x-value of shaded regions moving across rows). In early years, the invasion covered relatively little area and formed one contiguous shape, and sampling effort was dense (Figure 2, Table 1). Under these conditions, small α’ values were favored (Figure 3B). By 2023, however, the invaded area was discontinuous and irregularly shaped, and sampling density was lower. Under these conditions, higher α’ values resulted in better performance. Higher β values resulted in coarser preferred spatial scales (Figure 3B, compare x-values of the shaded region within columns). For the α- convex hull analysis, the optimal scale based on averaging across years ranged from α’ = 12 km with β = 1 to α’ = 100 km with β = 4.

For further analyses, we selected the optimal value of α’ = 35 km based on the pooled contingency table results across years with β = 2. Choosing a β = 2 in the F_β_ score emphasizes recall over precision, making this β value particularly useful for identifying boundaries that prioritize capturing all potential presences. Thus, a β = 2 is advantageous when it is critical to avoid underestimating the invaded range missing outlying populations.

### Satellite population spread analysis

We identified 153 distinct satellite populations over the course of *L. delicatula*’s invasion in the U.S. based on α-convex hulls drawn with α’ = 35 km. Of these, 46 persisted as distinct, disjointed invasion epicenters for at least 1 year without being subsumed into the primary invasion. However, not all satellite populations were sampled intensively enough to characterize year-to-year boundary displacement. Specifically, only 29 of the 1-year-old epicenters were sufficiently sampled, having at least 20 recorded absences within 5 km of presences in each year of observation (years 0 and 1). Overall, the epicenters exhibited slow initial spread, and in several cases the invaded range did not change between the year of first discovery and the next, despite continued sampling around the selected satellite populations. The lack of observed spread given existing sampling effort suggests any such spread was limited. Median boundary displacement distance for satellite populations in their first year ranged from 0 km to 10.6 km. In their second year, boundary displacement was larger, generally, with median displacement ranging from 0 km to 15.4 km. The only satellite population that persisted for 5 years or longer during the study period was the primary invasion originating in Berks County, PA. Boundary displacement around the primary invasion slowed in later years, with median boundary displacement falling from its peak value of 20.7 km when the invasion had been known for 3 years (in 2017) to 5.1, 7.5, and 4.3 km in its 7^th^, 8^th^, and 9^th^ years respectively.

## Discussion

Analysis of the well-documented record of *Lycorma delicatula*’s distribution over time in the United States revealed that the invasion’s spread rate has varied over time, generally following the slow-fast-slow pattern expected for biological invasions (Figure 2) with peak yearly boundary displacement of around 21 km/yr. The >900,000 presence and absence point records we aggregated allowed us to evaluate the influence of spatial scaling and employ cross validation to select an optimal spatial scale to estimate spread patterns (Figure 1, 3). Analysis with coarser scales resulted in faster spread rate estimates (Figure 2). Generally, finer scale parameters were favored in early years when sampling density was high and the distribution was compact and contiguous, while intermediate scaling parameters performed better in later years when sampling density was lower and the distribution discontinuous and irregular (Figure 3). In their first year of spread, isolated satellite populations typically spread short distances, with median boundary displacements ranging from 0 km to 10.6 km (Figure 4). Most satellite populations had median boundary displacement under 1.5 km in the first year after discovery. Together, these findings demonstrate the importance of considering temporal variation and spatial scaling when deriving spread pattern estimates based on geolocated point data and highlight the importance of targeting nascent satellite populations with management before their spread rates accelerate to the rapid rates observed around primary epicenters invasions.

**Figure 4:**
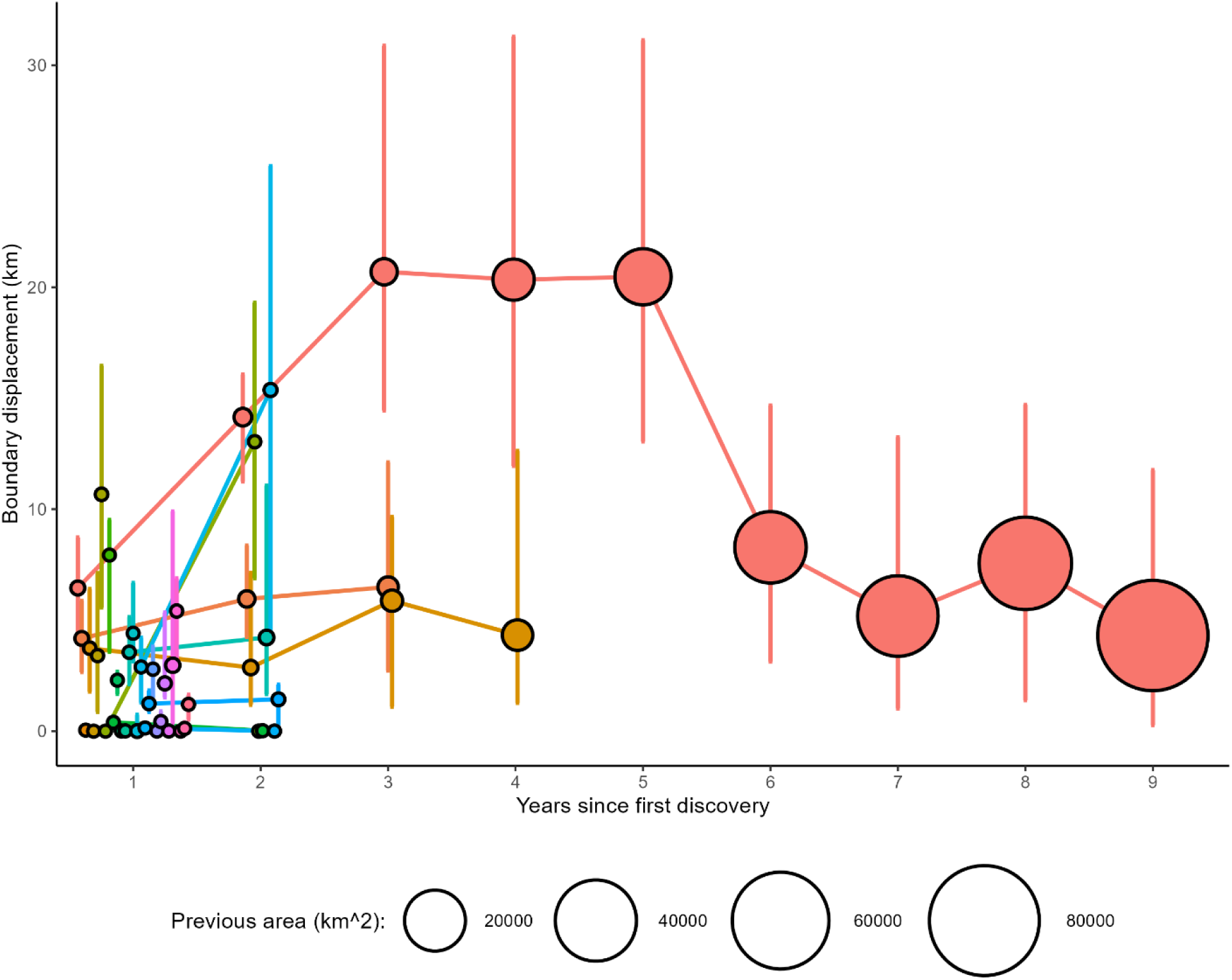
Spread rate varied with satellite population age. Distances between consecutive year boundaries of *L. delicatula* satellite populations were measured using α-convex hulls. Most epicenters, located near the primary invasion were subsumed within a few years. Points represent median boundary displacement, with bars showing the inter-quartile range. Epicenters are uniquely colored, and points are connected to track changes over time. In the first year after detection, most epicenters exhibited shorter spread distances compared to the >20 km annual spread observed for the primary invasion 3-5 years after its detection.

We observed rapid annual spread around the primary *L. delicatula* invasion during the years of peak spread, with boundaries moving a median distance of 20.7 (IQR: 14.5 – 30.9), 20.3 (IQR: 12.0 – 31.3), and 20.5 km (IQR: 13.1 – 31.1 km) in 2017, 2018, and 2019 respectively.

The spread of *L. delicatula* in the U.S. has been comparable to that of other introduced insect pests. The spongy moth (*Lymantria dispar*), for example, was estimated to spread at an average rate of 21 km yr^-1^ from 1966 to 1990 before the release of biological control agents (Liebhold et al., 1992). The Japanese beetle (*Popillia japonica*) was estimated to spread at 11.9 km/year for 1939-1951 (Allsopp, 1996). Finally, a median spread rate of 17.7 km/year (range 0.014 to 400 km/year) has been observed across 86 insect species introduced to the U.S. (Fahrner & Aukema, 2018).

For *L. delicatula,* future work should focus on explaining variation in spread patterns, based on land cover, transport flows, and environmental conditions (Andrew & Ustin, 2010; Tobin & Robinet, 2022). Prior analysis of *L. delicatula* spread rates found an association between human population density and *L. delicatula* spread, with denser human populations increasing the likelihood that counties were colonized by this pest (Cook et al. 2020). Models of *L. delicatula* spread incorporate the influence of dispersal pathways including rail, transport corridors and trade connections, which can potentially help individuals to move long distances and colonize new areas (Huron et al., 2022; Jones et al., 2021; Ladin et al., 2023). Further, *L. delicatula* population dynamics are highly temperature-dependent, influencing survival, reproduction, and development, and should be explicitly incorporated into future research aiming to explain and model spread patterns (Kreitman et al., 2021; Lewkiewicz et al., 2022, 2024).The spread of *L. delicatula* has involved repeated long-distance transport events (Figure 1)(Urban et al., 2021). Future research should focus on identifying property types that increase the likelihood of generating propagules capable of transport to better target management efforts. Additionally, understanding the types of properties that support the establishment of new satellite population epicenters could enhance surveillance strategies.

Cross-validation is a robust method for identifying an appropriate spatial scale for spread pattern analyses. Historically, data availability often constrained the choice of spatial scale; for instance, occurrence data documented at the county level effectively set the minimum viable resolution for analysis (Liang et al., 2020). Today, however, the widespread availability of geolocated point data—provided by citizen scientists contributing to online platforms and by trained surveyors using accurate and accessible GPS technology—offers finer resolution for mapping invader distributions (Crall et al., 2011; Nathan et al., 2022). Despite these advancements, researchers must still devise methods to delineate invaded from uninvaded areas, effectively locating the invasion front. The goal is to establish boundaries capable of accurately distinguishing additional presence and absence locations if further observations were made.

Cross-validation simulates this process by holding aside a subset of the data to evaluate how well boundaries drawn from the remaining data predict the withheld records. Future work should explore how different cross-validation strategies influence the selection of spatial scales for spread pattern analyses, particularly in the context of highly autocorrelated, heterogeneous, or fragmented landscapes (Dewhirst & Lutscher, 2009; Meyer et al., 2019). Investigating the impact of varying the proportion of data withheld and the spatial arrangement of withheld points could provide insights into the robustness and generalizability of boundary delineation methods (Roberts et al., 2017). Finally, extending cross-validation techniques to better account for temporal lags and spread rate variation could enhance predictions of invasion front progression over time, providing more reliable tools for management and surveillance (Aikio et al., 2010; Lampert, 2024; Neubert & Parker, 2004).

Comparing the performance of different spatial scales when delineating invaded ranges requires selecting an evaluation metric, with the F_β_ score being particularly useful because it encapsulates two key properties relevant to invasion biology: precision and recall. Precision is the proportion of true positives (correctly predicted presences) out of all predicted presences (true positives plus false positives). High precision minimizes false positives, ensuring resources are directed to genuinely invaded areas. Recall (or sensitivity) is the proportion of true positives out of all actual presences (true positives plus false negatives), capturing as much of the invasion range as possible. Depending on the species and available resources, stakeholders must balance precision and recall, prioritizing precision to avoid unnecessary interventions or recall ensuring critical invaded areas are not overlooked (Sofaer et al., 2019). The F_β_ metric allows for adjustment of the relative importance placed on each, providing flexibility in stakeholder preference. Alternative metrics, such as the commonly used Matthew’s Correlation Coefficient (MCC), do not provide the flexibility to adjust the relative importance of precision and recall, making them less suitable for invaded range delineation for surveys where finding true positives is critical.

The F_β_ metric originated in the field of information retrieval, where for a given query, the number of irrelevant texts is exceedingly large. In such cases, it is valuable to focus on precision and recall, which are unaffected by the number of correctly classified negative cases (Christen et al., 2024). Similarly, there are innumerable locations where the target pest does not occur, and identifying such locations is of little value. Instead, defining a range boundary that includes as many presences as possible (high recall) while excluding absences as much as possible (high precision) is desirable. Flexible weighting of these two goals is achieved by altering the β parameter in the F_β_ metric. We found that β values for the *L. delicatula* invasion of the US ranging from 1-3 generally produced clear peaks in optimal scaling parameters (cell edge length for grids and α for α-convex hulls, Figure 3). Higher values of β, reflecting increased importance placed on capturing all presence records over limiting the inclusion of absences, resulted in coarser preferred α values. Very high β values were unworkable, however, showing a broad range of α values that performed almost identically well, because once nearly all presences are being captured, including more absences has relatively little impact with high β values. We recommend the use of F_β_ as a tool to evaluate binary distribution maps of invasive species spread, allowing practitioners to select appropriate spatial scales in their analyses based on their particular assessment of the relative importance of recall and precision for their targeted pest. Future work should examine how the choice of β might be constrained by data properties, such as class imbalance. Similarly, other metrics that balance precision and recall should also be explored and contrasted to F_β_ (Li & Guo, 2021; Sofaer et al., 2019).

Our approach to locating *L. delicatula’*s invasion front was based on presence records made by state and federal agencies, and absences were used only to evaluate the boundaries drawn using these presences. This approach provides a highly detailed picture of the spread of this pest, but alternative methods may add new insights in the future. Public reports of *L. delicatula* presence provide a wealth of information, but are likely to include some number of false positive reports that may hamper analysis, although methods for estimating the false positive rate are available (Clare et al., 2019). Our analysis also did not account for imperfect detection, where observers may record an absence when the targeted organism is truly present (Berec et al., 2015). Future work to estimate detection rates (Valentin et al., 2020) may help to more completely incorporate absence records into future analyses of *L. delicatula* spread dynamics.

The spread of biological invasions is a dynamic process, and spread rates are expected to change over the course of the invasion (Shigesada et al., 1995). We found the expected pattern, with slow initial spread accelerating to reach a relatively constant rate of spread. Eventually, as invasions reach the margins of suitable habitat in the invaded range, their spread rate is expected to slow (Shigesada et al., 1995). The location of range limits and the timing when *L. delicatula* is expected to reach those limits are both uncertain, and subject to change with a warming and increasingly variable climate (Lewkiewicz et al., 2024; Zhao et al., 2024). Continued monitoring of *L. delicatula*’s occupancy patterns can provide useful information on ongoing spread, informing both managers who require distribution information to target treatments and ecologists who can learn about spread patterns as populations reach and exceed climatic range limits (Keena et al., 2023). Concerningly, multiple analyses indicate that the wine growing regions on the west coast of the United States are suitable for establishment by *L. delicatula* (Huron et al., 2022). Preventing spread to these locations remains an important goal.

We observed a slowdown in boundary displacement around the primary epicenter originating in Berks County, PA in the years 2020 through 2023. The causes for this pattern remain unclear. The invasion reached the eastern shoreline during this period. The invasion’s margin also reached outside Pennsylvania for the first time during this period, and differences may therefore be caused in part by different sampling plans used by different state agencies surveying this pest’s distribution (Figure 1). The COVID-19 pandemic and shutdown also affected survey efforts during these years, reducing the sampling effort even as the invasion’s border continued to grow (Table 1). The slowdown in spread might also be caused by biological factors. Climatic conditions might have been less favorable in these years than in 2018 when boundary expansion was at its most rapid. Because the invasion is relatively new, the sample size to make strong statements regarding climate’s impact on spread rates remains very low, however. Alternatively, accumulating natural enemies may slow population growth, thereby reducing spread (Johnson et al., 2023). It may also be true that more frequent establishment of small, isolated populations close to the margins of the primary invasion affected our measure of boundary displacement, which drew the shortest distance between one year’s boundary and any occupied area in the prior year, not just to the targeted population’s prior limit. With more small forerunner populations, drawn boundary displacement may be short, even if the main boundary moved a substantial distance.

*Lycorma delicatula’*s spread in its invaded range in the United States is ongoing.

Continuing establishment of far-flung outlying populations threatens to continue the rapid range expansion that we have observed in the years 2014 through 2023. Continued sampling effort is necessary to inform better mechanistic understanding of the ways in which this pest spreads. By looking at the history of spread, particularly the history of spread in the first few years after first detection, managers can derive useful projections for how far newly established populations that might yet be targeted for eradication are likely to spread. Slow initial spread should motivate managers to take early action and treat nascent satellite populations before their spread rate accelerates. Additionally, analyses of invasive spread patterns should carefully consider the scale used to derive estimates, as differences between scales can substantially alter estimated spread rates and the locations of invaded range boundaries. Effective surveying for *L. delicatula* and similar invasive species will require leveraging historical spread patterns, optimizing spatial scales for analysis, and incorporating cross-validation to improve range boundary accuracy. By combining these approaches with early intervention at nascent satellite populations, managers can mitigate and slow the long-term impacts of invasive species.

## Acknowledgements

For sharing data on *L. delicatula* occurrence, we thank the U.S. Department of Agriculture Spotted Lanternfly Working Group, the Pennsylvania Department of Agriculture, and the New York State Department of Agriculture and Markets, the Massachusetts Department of Agricultural Resources, and the North Carolina Dept. of Agriculture and Consumer Services. This work was funded by the United States Department of Agriculture (USDA) Animal and Plant Health Inspection Service Plant Protection and Quarantine under agreements AP22PPQS&T00C146, AP22PPQS&T00C097, AP23PPQS&T00C090, and AP24PPQS&T00C136; the USDA National Institute of Food and Agriculture Specialty Crop Research Initiative under the coordinated agricultural project award 2019-51181-30014; the USDA Tactical Sciences for Agricultural Biosecurity under the project award 2022-68013- 37139; the Pennsylvania Department of Agriculture under agreements C94000833 and C940001671.

## Author Contributions

All authors conceived of the study and contributed to gathering data. JK carried out analyses and wrote the first draft of the manuscript. All authors contributed to revising the manuscript and approved the final draft.

## Conflict of Interest Statement

The authors declare no conflict of interest

## Appendix S1

**Table S1:** Regression results, with invaded area determined based on intersection of established points with grids of various scales, fit using simple linear regression. Rows for which simple linear regression had better fit than non-linear logistic regression (see table S2) are shown in bold.

| edge | AICc | F | numerator<br>df | denominator<br>df | p-value | intercept | intercept<br>S.E. | slope | slope S.E. |
| --- | --- | --- | --- | --- | --- | --- | --- | --- | --- |
| 1000 | 61.46 | 278.2 | 1 | 8 | 1.69E-07 | -13109.7 | 787.7 | 6.51 | 0.39 |
| 2000 | 69.49 | 310.1 | 1 | 8 | 1.11E-07 | -20682.1 | 1177.1 | 10.27 | 0.58 |
| 4000 | 72.99 | 453.5 | 1 | 8 | 2.49E-08 | -29796.9 | 1402.3 | 14.79 | 0.69 |
| **6000** | **72.90** | **644.9** | **1** | **8** | **6.19E-09** | **-35383.2** | **1396.3** | **17.57** | **0.69** |
| **8000** | **72.88** | **787.4** | **1** | **8** | **2.81E-09** | **-39049.8** | **1394.6** | **19.39** | **0.69** |
| **10000** | **72.48** | **968.1** | **1** | **8** | **1.24E-09** | **-42444.9** | **1367.0** | **21.07** | **0.68** |
| **12000** | **73.58** | **956.0** | **1** | **8** | **1.30E-09** | **-44564.3** | **1444.3** | **22.12** | **0.72** |
| 14000 | 78.17 | 683.1 | 1 | 8 | 4.93E-09 | -47386.5 | 1816.9 | 23.52 | 0.90 |
| 16000 | 77.60 | 771.0 | 1 | 8 | 3.05E-09 | -48932.3 | 1765.9 | 24.29 | 0.87 |
| 18000 | 80.49 | 621.2 | 1 | 8 | 7.18E-09 | -50751.9 | 2040.6 | 25.20 | 1.01 |
| 20000 | 82.42 | 561.1 | 1 | 8 | 1.07E-08 | -53125.3 | 2247.3 | 26.37 | 1.11 |
| 25000 | 83.81 | 543.9 | 1 | 8 | 1.21E-08 | -56055.8 | 2408.6 | 27.83 | 1.19 |
| 30000 | 87.18 | 435.2 | 1 | 8 | 2.92E-08 | -59330.8 | 2850.1 | 29.46 | 1.41 |
| 35000 | 89.02 | 411.1 | 1 | 8 | 3.66E-08 | -63224.5 | 3124.8 | 31.39 | 1.55 |
| 40000 | 91.88 | 337.4 | 1 | 8 | 7.94E-08 | -66111.9 | 3606.6 | 32.82 | 1.79 |
| 45000 | 93.87 | 297.3 | 1 | 8 | 1.30E-07 | -68547.1 | 3983.8 | 34.03 | 1.97 |
| 50000 | 93.49 | 321.9 | 1 | 8 | 9.55E-08 | -69970.3 | 3908.5 | 34.74 | 1.94 |
| 75000 | 102.08 | 181.2 | 1 | 8 | 8.89E-07 | -80685.3 | 6006.3 | 40.06 | 2.98 |
| 100000 | 103.60 | 183.8 | 1 | 8 | 8.41E-07 | -87645.0 | 6479.0 | 43.52 | 3.21 |

**Table S2:** Regression results, with invaded area determined based on intersection of established points with grids of various scales, fit using nonlinear, logistic regression. Rows for which this non-linear regression had better fit than the corresponding simple linear regression (see table S1) are bolded.

| edge | AICc | F | numerator<br>df | denominator<br>df | p-value | asymptote | asymptote<br>S.E. | midpoint | midpoint<br>S.E. | logistic<br>slope | logistic slope<br>S.E. |
| --- | --- | --- | --- | --- | --- | --- | --- | --- | --- | --- | --- |
| **1000** | **55.67** | **403.6** | **2** | **7** | **5.89E-08** | **56.49** | **2.30** | **2018.25** | **0.20** | **1.47** | **0.15** |
| **2000** | **63.64** | **451.5** | **2** | **7** | **3.99E-08** | **88.43** | **3.52** | **2018.35** | **0.20** | **1.46** | **0.14** |
| **4000** | **70.23** | **481.3** | **2** | **7** | **3.20E-08** | **130.08** | **5.67** | **2018.56** | **0.22** | **1.56** | **0.15** |
| 6000 | 74.43 | 443.3 | 2 | 7 | 4.25E-08 | 160.50 | 8.56 | 2018.82 | 0.27 | 1.70 | 0.17 |
| 8000 | 75.00 | 509.6 | 2 | 7 | 2.62E-08 | 183.10 | 10.19 | 2019.02 | 0.28 | 1.79 | 0.17 |
| 10000 | 76.67 | 508.2 | 2 | 7 | 2.65E-08 | 202.05 | 12.11 | 2019.16 | 0.31 | 1.83 | 0.18 |
| 12000 | 77.32 | 525.0 | 2 | 7 | 2.36E-08 | 217.40 | 13.67 | 2019.27 | 0.32 | 1.90 | 0.18 |
| **14000** | **77.48** | **587.0** | **2** | **7** | **1.60E-08** | **242.32** | **16.70** | **2019.59** | **0.36** | **1.98** | **0.19** |
| **16000** | **74.24** | **865.8** | **2** | **7** | **4.14E-09** | **250.21** | **13.93** | **2019.54** | **0.29** | **1.99** | **0.15** |
| **18000** | **75.92** | **789.2** | **2** | **7** | **5.72E-09** | **266.68** | **16.93** | **2019.73** | **0.33** | **2.04** | **0.17** |
| **20000** | **75.12** | **937.8** | **2** | **7** | **3.13E-09** | **264.46** | **13.37** | **2019.49** | **0.26** | **1.90** | **0.14** |
| **25000** | **69.81** | **1781.3** | **2** | **7** | **3.34E-10** | **292.53** | **11.83** | **2019.65** | **0.21** | **2.01** | **0.11** |
| **30000** | **72.09** | **1593.9** | **2** | **7** | **4.92E-10** | **328.25** | **16.26** | **2019.95** | **0.26** | **2.13** | **0.12** |
| **35000** | **76.49** | **1166.0** | **2** | **7** | **1.47E-09** | **327.20** | **16.01** | **2019.64** | **0.25** | **1.97** | **0.13** |
| **40000** | **80.71** | **838.3** | **2** | **7** | **4.63E-09** | **352.18** | **21.90** | **2019.79** | **0.32** | **2.02** | **0.16** |
| **45000** | **70.31** | **2563.8** | **2** | **7** | **9.36E-11** | **379.54** | **14.60** | **2019.96** | **0.20** | **2.09** | **0.09** |
| **50000** | **76.18** | **1481.3** | **2** | **7** | **6.36E-10** | **399.63** | **21.49** | **2020.03** | **0.29** | **2.19** | **0.13** |
| **75000** | **88.78** | **567.3** | **2** | **7** | **1.81E-08** | **473.16** | **43.12** | **2020.20** | **0.48** | **2.18** | **0.21** |
| **100000** | **96.45** | **309.1** | **2** | **7** | **1.49E-07** | **578.44** | **90.40** | **2020.62** | **0.89** | **2.51** | **0.34** |

**Table S3:** Regression results, with invaded area determined based on the α-convex hull around established points, fit using simple linear regression. Rows for which simple linear regression had better fit than non-linear logistic regression (see table S4) are shown in bold.

| $\alpha$ | $\alpha'$ | AICc | F | numerator df | denominator df | p-value | intercept | intercept S.E. | slope | slope S.E. |
| --- | --- | --- | --- | --- | --- | --- | --- | --- | --- | --- |
| 564 | 1000 | 38.98 | 170.5 | 1 | 8 | 1.12E-06 | -3334.0 | 256.0 | 1.66 | 0.13 |
| 1128 | 2000 | 54.99 | 175.8 | 1 | 8 | 1.00E-06 | -7538.1 | 570.0 | 3.74 | 0.28 |
| 2257 | 4000 | 71.22 | 141.9 | 1 | 8 | 2.27E-06 | -15251.6 | 1283.3 | 7.57 | 0.64 |
| 3385 | 6000 | 78.70 | 153.1 | 1 | 8 | 1.70E-06 | -23036.8 | 1865.8 | 11.44 | 0.92 |
| 4514 | 8000 | 80.31 | 176.8 | 1 | 8 | 9.78E-07 | -26822.5 | 2021.8 | 13.32 | 1.00 |
| 5642 | 10000 | 77.35 | 285.3 | 1 | 8 | 1.53E-07 | -29395.3 | 1744.2 | 14.59 | 0.86 |
| 6770 | 12000 | 75.66 | 382.6 | 1 | 8 | 4.85E-08 | -31271.3 | 1602.1 | 15.52 | 0.79 |
| **7899** | **14000** | **74.06** | **504.2** | **1** | **8** | **1.64E-08** | **-33144.1** | **1479.2** | **16.45** | **0.73** |
| **9027** | **16000** | **72.39** | **642.5** | **1** | **8** | **6.29E-09** | **-34413.2** | **1360.5** | **17.08** | **0.67** |
| **10155** | **18000** | **72.13** | **695.5** | **1** | **8** | **4.59E-09** | **-35355.8** | **1343.4** | **17.55** | **0.67** |
| **11284** | **20000** | **70.76** | **840.5** | **1** | **8** | **2.17E-09** | **-36290.0** | **1254.4** | **18.02** | **0.62** |
| **14105** | **25000** | **71.67** | **876.5** | **1** | **8** | **1.84E-09** | **-38774.6** | **1312.4** | **19.25** | **0.65** |
| **16926** | **30000** | **72.71** | **848.9** | **1** | **8** | **2.09E-09** | **-40210.6** | **1382.9** | **19.96** | **0.69** |
| **19747** | **35000** | **75.49** | **716.6** | **1** | **8** | **4.08E-09** | **-42446.6** | **1588.8** | **21.07** | **0.79** |
| **22568** | **40000** | **78.28** | **613.0** | **1** | **8** | **7.57E-09** | **-45137.3** | **1826.6** | **22.41** | **0.90** |
| **25389** | **45000** | **79.13** | **604.0** | **1** | **8** | **8.03E-09** | **-46755.9** | **1906.2** | **23.21** | **0.94** |
| 28209 | 50000 | 80.91 | 536.0 | 1 | 8 | 1.29E-08 | -48148.4 | 2083.7 | 23.90 | 1.03 |
| 42314 | 75000 | 84.04 | 465.6 | 1 | 8 | 2.24E-08 | -52479.4 | 2436.6 | 26.05 | 1.21 |
| 56419 | 100000 | 88.54 | 359.7 | 1 | 8 | 6.18E-08 | -57767.5 | 3051.5 | 28.67 | 1.51 |

**Table S4:** Regression results, with invaded area determined based on the α-convex hull around established points, fit using nonlinear, logistic regression. Rows for which this non-linear regression had better fit than the corresponding simple linear regression (see table S3) are bolded.

| $\alpha$ | $\alpha'$ | AICc | F | numerator<br>df | denominator<br>df | p-value | intercept | asymptote | asymptote<br>S.E. | midpoint | midpoint<br>S.E. | rate | rate<br>S.E. |
| --- | --- | --- | --- | --- | --- | --- | --- | --- | --- | --- | --- | --- | --- |
| **564** | **1000** | **37.73** | **157.8** | **2** | **7** | **1.51E-06** | **NA** | **15.35** | **0.85** | **2017.43** | **0.31** | **1.61** | **0.25** |
| **1128** | **2000** | **50.55** | **224.7** | **2** | **7** | **4.47E-07** | **NA** | **33.44** | **1.65** | **2017.89** | **0.26** | **1.50** | **0.20** |
| **2257** | **4000** | **64.16** | **238.6** | **2** | **7** | **3.63E-07** | **NA** | **62.87** | **2.75** | **2018.02** | **0.21** | **1.24** | **0.17** |
| **3385** | **6000** | **67.77** | **379.7** | **2** | **7** | **7.28E-08** | **NA** | **91.10** | **3.03** | **2018.04** | **0.16** | **1.12** | **0.13** |
| **4514** | **8000** | **69.46** | **432.2** | **2** | **7** | **4.65E-08** | **NA** | **106.13** | **3.43** | **2018.10** | **0.15** | **1.16** | **0.12** |
| **5642** | **10000** | **70.35** | **467.3** | **2** | **7** | **3.54E-08** | **NA** | **118.35** | **4.07** | **2018.20** | **0.16** | **1.28** | **0.13** |
| **6770** | **12000** | **75.02** | **328.1** | **2** | **7** | **1.21E-07** | **NA** | **128.58** | **5.85** | **2018.33** | **0.22** | **1.39** | **0.17** |
| 7899 | 14000 | 77.77 | 278.2 | 2 | 7 | 2.14E-07 | NA | 139.15 | 7.51 | 2018.44 | 0.26 | 1.49 | 0.19 |
| 9027 | 16000 | 78.68 | 272.9 | 2 | 7 | 2.29E-07 | NA | 148.03 | 8.85 | 2018.59 | 0.30 | 1.58 | 0.21 |
| 10155 | 18000 | 79.21 | 272.7 | 2 | 7 | 2.29E-07 | NA | 153.87 | 9.65 | 2018.67 | 0.31 | 1.61 | 0.21 |
| 11284 | 20000 | 79.35 | 283.1 | 2 | 7 | 2.01E-07 | NA | 160.68 | 10.54 | 2018.78 | 0.33 | 1.67 | 0.22 |
| 14105 | 25000 | 81.80 | 252.5 | 2 | 7 | 2.99E-07 | NA | 181.22 | 15.39 | 2019.17 | 0.43 | 1.81 | 0.25 |
| 16926 | 30000 | 82.27 | 259.2 | 2 | 7 | 2.73E-07 | NA | 193.11 | 17.73 | 2019.35 | 0.47 | 1.87 | 0.26 |
| 19747 | 35000 | 83.23 | 262.7 | 2 | 7 | 2.61E-07 | NA | 211.82 | 21.94 | 2019.63 | 0.53 | 1.94 | 0.28 |
| 22568 | 40000 | 81.47 | 356.3 | 2 | 7 | 9.08E-08 | NA | 229.32 | 21.74 | 2019.81 | 0.48 | 1.94 | 0.24 |
| 25389 | 45000 | 79.26 | 478.1 | 2 | 7 | 3.27E-08 | NA | 233.15 | 18.18 | 2019.74 | 0.39 | 1.89 | 0.20 |
| **28209** | **50000** | **77.90** | **582.5** | **2** | **7** | **1.65E-08** | **NA** | **238.51** | **16.58** | **2019.74** | **0.34** | **1.85** | **0.18** |
| **42314** | **75000** | **79.53** | **589.1** | **2** | **7** | **1.58E-08** | **NA** | **256.32** | **17.27** | **2019.73** | **0.32** | **1.80** | **0.17** |
| **56419** | **100000** | **79.18** | **744.3** | **2** | **7** | **7.02E-09** | **NA** | **286.77** | **18.32** | **2019.91** | **0.30** | **1.80** | **0.16** |

**Figure S1:**
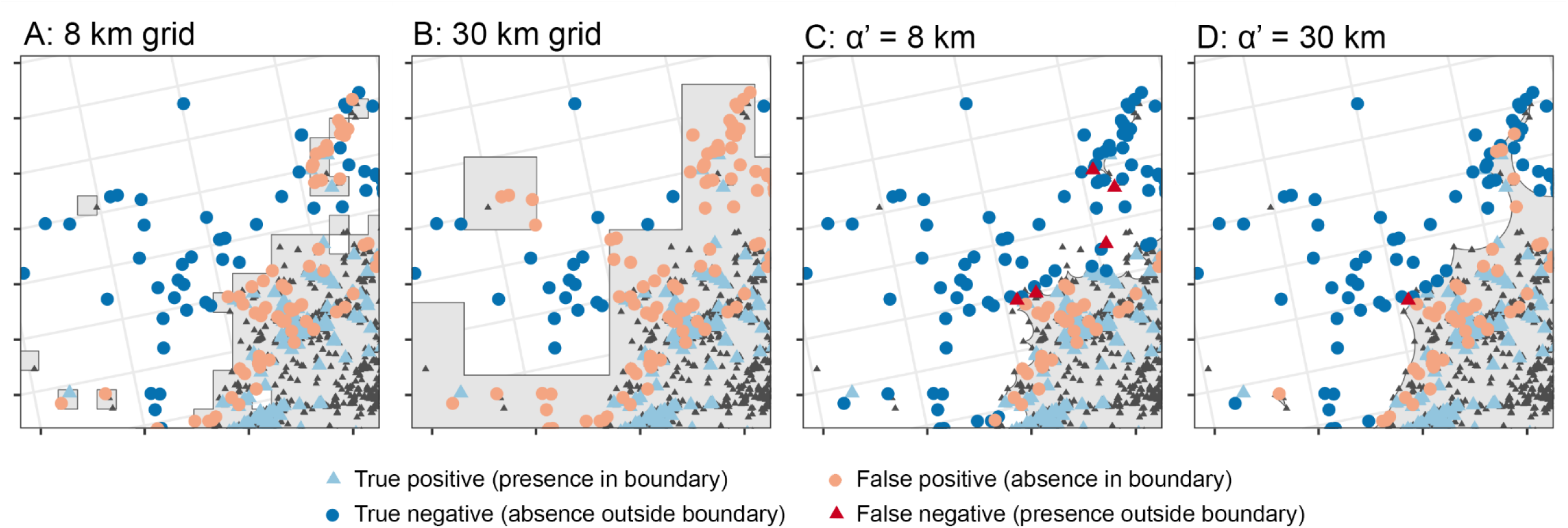
Cross validation allows for the selection of a best performing spatial scale for analysis. Examples of differing grid scales are shown in panels A and B, and α-convex hulls with differing α’ values in C and D. In these panels, 4/5 of the data were used to derive the boundary (showing as small grey points) and the remaining 1/5 to evaluate the boundary, (points colored according to their status).

## Notes

https://github.com/ieco-lab/lydemapr

### Competing Interest Statement

The authors have declared no competing interest.

